# Target validation uncouples TSPO from 19-Atriol-mediated inhibition of steroidogenesis and reveals true enzymatic targets

**DOI:** 10.1101/2025.09.25.678677

**Authors:** Amy H. Zhao, Prasanthi P. Koganti, Mingxing Qian, Anthony Garcia, Patrick O’Day, Richard J. Auchus, Douglas F. Covey, Vimal Selvaraj

## Abstract

The mitochondrial translocator protein (TSPO) was once proposed to mediate mitochondrial cholesterol import for steroid hormone biosynthesis, but genetic deletion studies in multiple models have refuted this role. Nevertheless, the idea that pharmacological ligands of TSPO can modulate steroid output continues to be invoked. One such compound, 19-Atriol (androst-5-ene-3β,17β,19-triol), was reported to inhibit progesterone synthesis via TSPO binding in MA-10 Leydig cells. To evaluate this proposed mechanism, we used CRISPR/Cas9-generated *Tspo*-deleted MA-10 cells to study 19-Atriol activity. We found that 19-Atriol inhibited Bt_2_-cAMP-stimulated steroid output independent of TSPO expression; it acted as a competitive inhibitor of 3β-hydroxysteroid dehydrogenase (3β-HSD), blocking the conversion of pregnenolone to progesterone. Mass spectrometry revealed that 19-Atriol is also a substrate for 3β-HSD, yielding 19-hydroxytestosterone (19-OHT), which itself inhibits 3β-HSD activity. In addition to this effect, both 19-Atriol and 19-OHT decreased cholesterol-to-pregnenolone conversion during stimulation. Partial inhibition of 22R-hydroxycholesterol metabolism by CYP11A1 was observed with 19-Atriol, but not 19-OHT, suggesting direct or indirect effects on this upstream step, potentially involving the steroidogenic acute regulatory protein (STAR). These findings decisively exclude TSPO as a functional mediator of 19-Atriol activity and instead identify direct enzymatic targets within the *de novo* steroidogenic pathway. By resolving a key mechanistic misattribution, this study underscores the importance of rigorous target validation, particularly for compounds previously assumed to act via TSPO.

## Introduction

Initially described as a peripheral benzodiazepine receptor distinct from the central benzodiazepine-binding GABA_A_ receptor, the precise function of the mitochondrial translocator protein (TSPO) has remained elusive (1, 2). Located in the outer mitochondrial membrane (3), TSPO first drew attention when certain small molecule binding compounds were shown to transiently increase steroid production (4–6). This observation led to the proposal that TSPO participates in mitochondrial cholesterol import (7), a process long regarded as the rate-limiting step in steroid hormone biosynthesis (8). This proposed role, amplified by substantial pharmacological interest in TSPO as a diagnostic and therapeutic target in neuroinflammation and neuropsychiatric disorders, fueled numerous studies that emphasized putative benefits while leaving the underlying mechanism unresolved in both model systems and humans (9–12). However, the development of multiple *Tspo* gene-knockout models, *in vivo* and *in vitro*, has refuted an essential role for TSPO in steroidogenesis (13–18) [reviewed in (19, 20)]. Moreover, genetic studies have established that that steroidogenic acute regulatory protein (STAR) as a critical mediator of mitochondrial cholesterol import (21), via an independent mechanism that does not involve TSPO (22). Consequently, the mechanisms with which the presumed TSPO binding drugs modulate steroidogenesis are unknown.

TSPO deficiency does not affect steroid hormone synthesis, but rather produces notable effects on lipid metabolism (23–25). Consistent with this finding, TSPO expression is not confined to steroidogenic tissues; higher expression correlates with lipid-enriched cells and with tissues active in lipid metabolism such as brown and white adipose tissue, harderian gland and lungs (24, 26, 27). Nevertheless, in the absence of a clear functional definition, pharmacological interpretations of TSPO-mediated effects have remained disparate and continue to invoke a core role in steroidogenesis (28, 29). A prominent example is the prototypical TSPO-binding drug PK11195 [N-butan-2-yl-1-(2-chlorophenyl)-N-methylisoquinoline-3-carboxamide], long considered an agonist and frequently cited as evidence for TSPO’s steroidogenic function (4–6), despite demonstrations in *Tspo*-deleted MA-10 Leydig cells (MA-10:*Tspo*^*Δ/Δ*^ cells) that PK11195’s steroidogenic effects are off-target and independent of TSPO (15). Beyond steroidogenesis, disparate cellular responses have been ascribed to TSPO-binding drugs, including programmed cell death (30), mitochondria-nuclear crosstalk (31), calcium homeostasis (32), ATP production (33), and generation and regulation of reactive oxygen species (34–36) further complicating efforts to define TSPO’s physiological role.

In a prior report, the steroid derivative 19-Atriol [androst-5-ene-3β,17β,19-triol] was reported to specifically bind TSPO and inhibit steroidogenesis in MA-10 Leydig cells (37). Binding to the cholesterol recognition amino acid consensus (CRAC) motif was proposed, implying competition with cholesterol for the same site on TSPO and thereby reducing mitochondrial cholesterol import (37). Subsequently, structure-activity relationships were further explored by steroid hydroxylation, offering a mechanism of specificity for TSPO inhibition for hydroxylation at C19 (such as seen in 19-Atriol) compared to others (hydroxylation at C4, C7 and C11) (38). *In vivo* studies administering 19-Atriol in rats also reported inhibition of Leydig cell testosterone production (39). Together, these reports positioned 19-Atriol as a putative TSPO antagonist and were used to support the broader model of TSPO involvement in steroidogenesis. However, because genetic studies have now excluded TSPO from an essential role in cholesterol import (13–18), the mechanism by which 19-Atriol influences steroid hormone synthesis remains unresolved. Here, we reevaluate the pharmacology of 19-Atriol in defined genetic systems to test whether its steroidogenic effects are mediated by TSPO or through alternative pathways.

## Results

### 19-Atriol inhibits progesterone synthesis independently of TSPO

To determine whether the previously reported inhibitory effect of 19-Atriol on steroidogenesis (37), was dependent on TSPO, we compared progesterone (P4) production in wild-type MA-10 Leydig cells and isogenic TSPO-null MA-10:*Tspo*^*Δ/Δ*^ cells. A schematic overview of the steroidogenic steps under consideration is shown in (Fig. 1A). Upon stimulation with dibutyryl cAMP (Bt_2_cAMP), both wild-type MA-10 and MA-10:*Tspo*^*Δ/Δ*^ cells exhibited a dose-dependent decrease in P4 synthesis when treated with 19-Atriol (Fig. 1B). The potency and maximal extent of inhibition were comparable between genotypes, demonstrating that TSPO is not required for 19-Atriol’s pharmacological activity.

**Figure 1.**
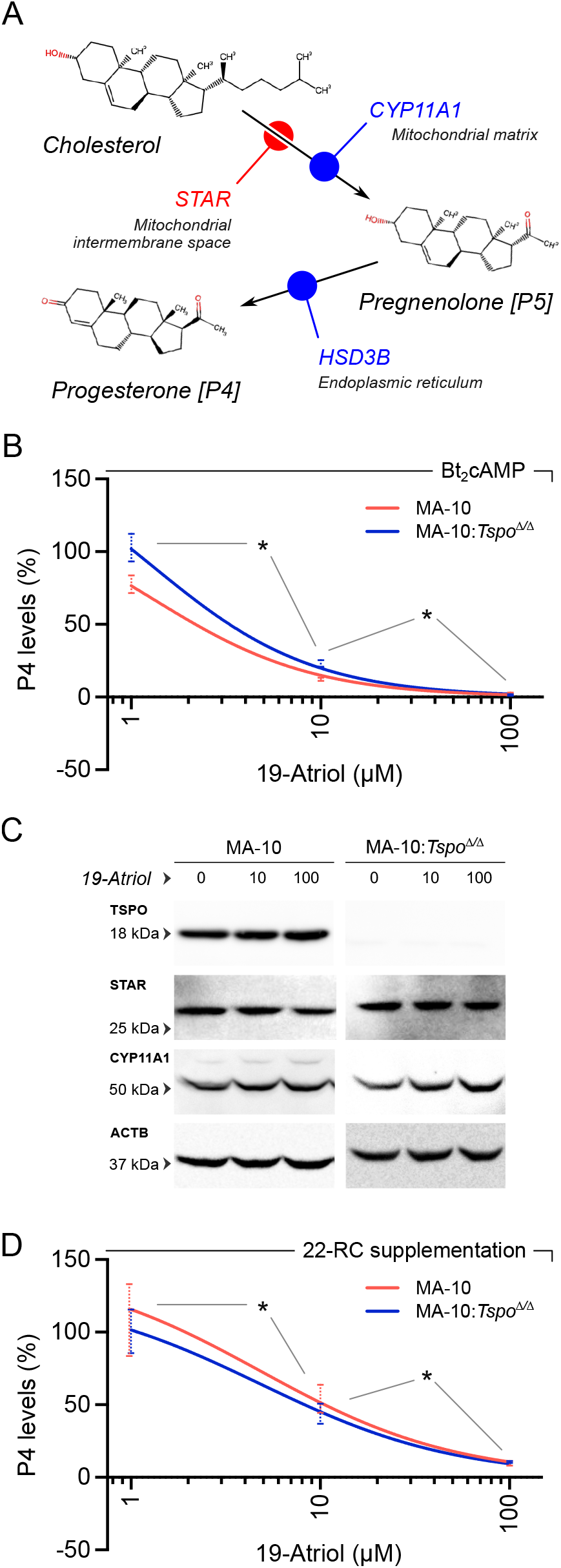
19-Atriol inhibits progesterone biosynthesis independently of TSPO and acts downstream of cholesterol import. **(A)** Schematic of the initial steps in the steroidogenic pathway: (i) Mitochondrial cholesterol import is mediated by the intermembrane space shuttle STAR. (ii) Cholesterol is converted to pregnenolone/P5 by CYP11A1 in the mitochondrial matrix. (iii) P5 is further converted to progesterone/P4 by 3β-hydroxysteroid dehydrogenase (HSD3B) at the endoplasmic reticulum. **(B)** Dose-response analysis of progesterone production in wild-type MA-10 and TSPO-null MA-10:*Tspo*^*Δ/Δ*^ cells stimulated with Bt_2_cAMP. Increasing concentrations of 19-Atriol reduced P4 output in both genotypes (*p<0.05) with comparable potency and maximal inhibition, indicating that its pharmacological action is independent of TSPO. **(C)** Immunoblot analysis of STAR and CYP11A1 expression following treatment with 19-Atriol (0,10 and 100 μM). TSPO was absent in MA-10:*Tspo*^*Δ/Δ*^ cells, validating the knockout. Neither STAR nor CYP11A1 protein abundance was altered by 19-Atriol in either genotype (ACTB served as a loading control). **(D)** Dose-response analysis of progesterone production in wild-type MA-10 and TSPO-null MA-10:*Tspo*^*Δ/Δ*^ cells with supplementation of 22R-HC in the absence of stimulation. 19-Atriol continued to inhibit P4 production in both genotypes (*p<0.05), indicating that the observed inhibitory effect occurs downstream of cholesterol import.

We next assessed whether 19-Atriol altered the abundance of proteins critical for the early steroidogenic pathway. Immunoblotting confirmed the absence of TSPO in MA-10:*Tspo*^*Δ/Δ*^ cells and revealed no detectable changes in STAR or CYP11A1 expression following treatment with 19-Atriol at concentrations up to 100 μM in either genotype (Fig. 1C). Thus, the observed inhibition was not attributable to altered expression of key steroidogenic proteins.

To localize the site of action within the pathway, we bypassed cholesterol import by supplementing cultures with the CYP11A1 intermediate 22(*R*)-hydroxycholesterol (22R-HC). Even under these conditions, 19-Atriol significantly reduced P4 output in both wild-type MA-10 and MA-10:*Tspo*^*Δ/Δ*^ cells, indicating that its inhibitory effect occurs downstream of cholesterol transport and side-chain cleavage (Fig. 1D). These results suggested that 19-Atriol acts on the enzymatic step converting pregnenolone (P5) to P4, prompting the need to directly evaluate 3β-hydroxysteroid dehydrogenase (HSD3B) activity.

### 19-Atriol inhibits the HSD3B-catalyzed conversion of pregnenolone to progesterone

To directly assess whether 19-Atriol acts on the step catalyzed by HSD3B, we supplemented cultures with P5 and monitored its conversion to P4. Addition of P5 (1-50 μM) produced a concentration-dependent increase in P4 accumulation in MA-10 cells (Fig. 2A), confirming efficient utilization of exogenous substrate and establishing a linear range suitable for inhibition studies. To validate assay specificity, we tested the well-characterized HSD3B inhibitor trilostane (40), which abolished P4 formation from P5 in both wild-type MA-10 and TSPO-null MA-10:*Tspo*^*Δ/Δ*^ cells (Fig. 2B). These results also verified that measured P4 production was strictly dependent on HSD3B activity and independent of TSPO.

**Figure 2.**
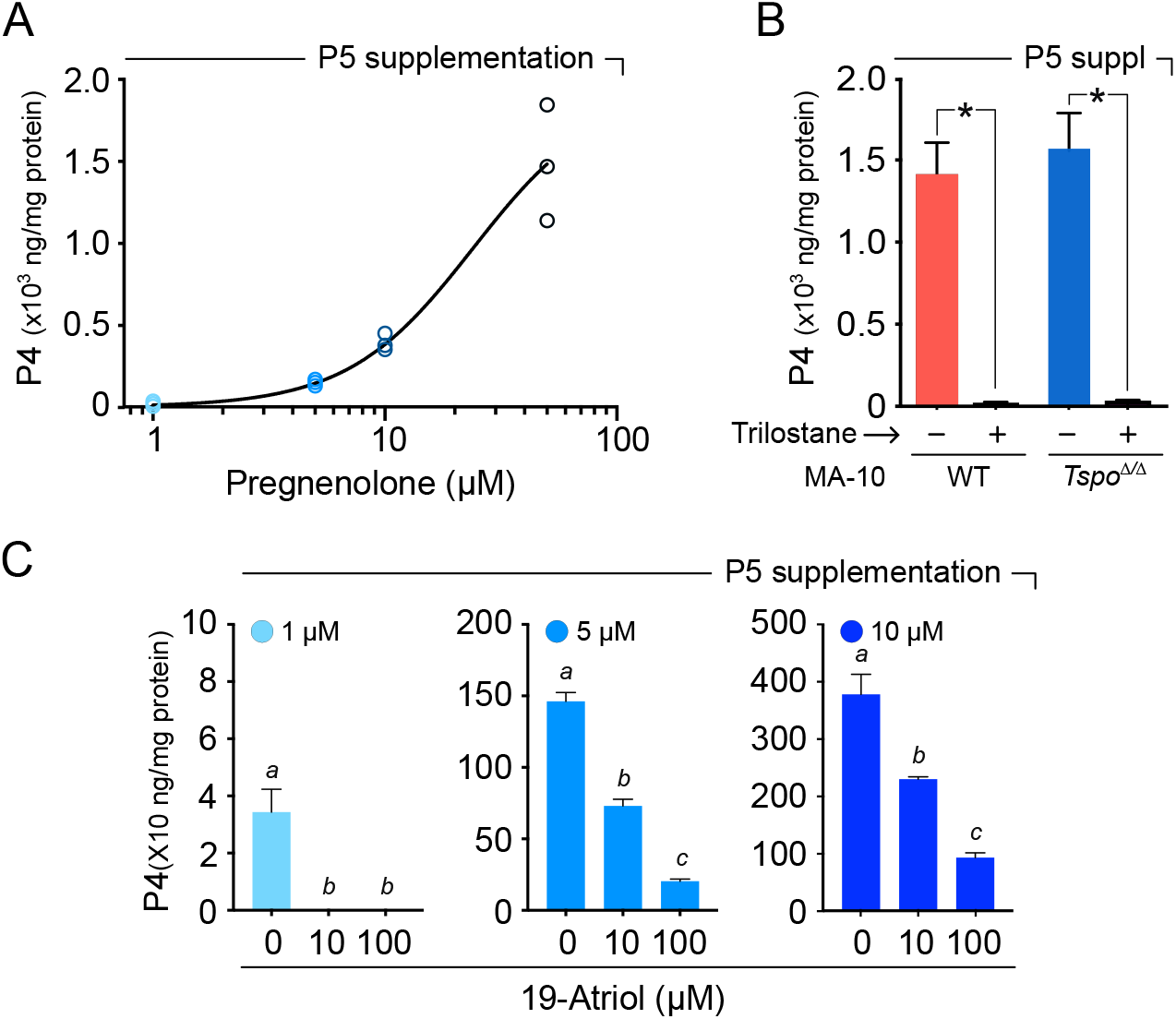
19-Atriol inhibits the HSD3B-catalyzed conversion of P5 to P4. **(A)** Titration of P5 in unstimulated MA-10 cell cultures demonstrated efficient conversion to P4. Increasing P5 (1-50 μM) produced a monotonic rise in P4 accumulation (normalized to total protein), establishing assay linearity and the dynamic range used for inhibitor testing. **(B)** Pharmacologic validation of the assay using the HSD3B inhibitor trilostane abolished P4 formation from exogenous P5 in both wild-type (WT) and TSPO-null MA-10:*Tspo*^*Δ/Δ*^ cells (*p<0.05). confirming that measured P4 derives from HSD3B activity and that this step does not require TSPO. **(C)** 19-Atriol inhibits P5 to P4 bioconversion in MA-10 cells. At three input P5 levels (1, 5, and 10 μM), 19-Atriol (0, 10, 100 μM) reduced P4 production in a dose-dependent manner. Distinct letters above bars denote statistically significant differences within each P5 condition (p<0.05).

When tested as an inhibitor, 19-Atriol significantly reduced the conversion of P5 to P4 in a dose-dependent manner (Fig. 2C). At 10 or 100 µM, 19-Atriol decreased or nearly abolished P4 output, respectively, for all three input concentrations of P5 (1, 5, and 10 μM). Statistical comparisons confirmed significant differences between treatment groups within each substrate condition (p < 0.05). These findings establish that 19-Atriol functions as an inhibitor of HSD3B.

### 19-Atriol is metabolized by HSD3B to 19-OHT, which also inhibits progesterone synthesis

Given the structural similarity to other 3β-hydroxy-Δ^5^-steroid substrates, we investigated whether 19-Atriol is a substrate for HSD3B. Mass spectrometry analysis demonstrated that treatment with 19-Atriol led to the formation of 19-hydroxytestosterone (19-OHT), both under basal and Bt_2_cAMP-stimulated conditions, consistent with metabolism by HSD3B (Fig. 3A). Accumulation of 19-OHT was concentration-dependent and saturable, indicating that 19-Atriol is enzymatically converted and providing a mechanistic basis for competition with endogenous substrates.

**Figure 3.**
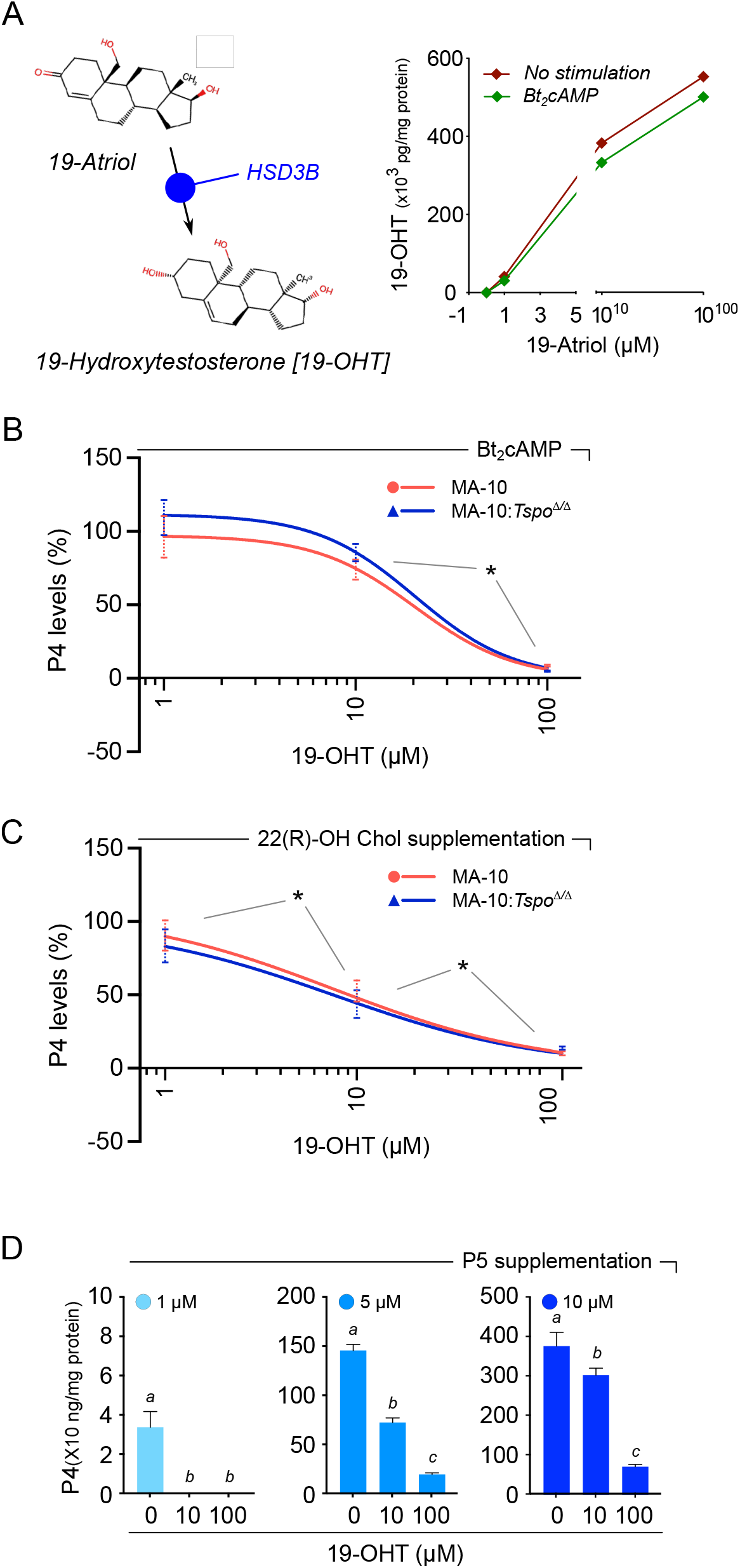
19-Atriol is metabolized by HSD3B to 19-OHT, and 19-OHT also inhibits progesterone synthesis. **(A)** Reaction scheme showing 19-Atriol is converted by 3β-hydroxysteroid dehydrogenase (HSD3B) to 19-hydroxytestosterone (19-OHT). LC–MS/MS showed dose-dependent accumulation of 19-OHT with increasing 19-Atriol (1–100 μM) in MA-10 cells under basal conditions and with Bt_2_cAMP stimulation, indicating that 19-Atriol serves as an HSD3B substrate. **(B)** Similar to 19-Atriol, 19-OHT decreased P4 output in a dose-dependent manner in wild-type MA-10 and TSPO-null MA-10:*Tspo*^*Δ/Δ*^ cells with comparable potency and maximal effect, with significant differences between the indicated concentrations (*p<0.05). **(C)** With 22R-HC supplementation, 19-OHT still reduced P4 production in both genotypes in a dose-dependent fashion similar to 19-Atriol, with significant differences between the indicated concentrations (*p<0.05). **(D)** In P5-supplemented assays, 19-OHT (0, 10, 100 μM) suppressed P4 formation at each P5 input (1, 5, and 10 μM). Distinct letters above bars indicate statistically significant differences within each P5 condition (p < 0.05).

We next tested the functional effects of 19-OHT on steroidogenesis. In Bt_2_cAMP-stimulated MA-10 cells, 19-OHT significantly reduced P4 output in a dose-dependent manner, with comparable inhibition in both wild-type MA-10 and TSPO-null MA-10:*Tspo*^*Δ/Δ*^ cells (Fig. 3B). Similarly, when the mitochondrial step was bypassed by supplementing with 22R-HC, 19-OHT continued to suppress P4 synthesis in both genotypes (Fig. 3C). These findings confirm that the inhibitory activity of 19-OHT, like that of 19-Atriol, is independent of TSPO.

Finally, direct substrate-product competition assays with P5 demonstrated that increasing concentrations of 19-Atriol reduced the conversion of P5 to P4 in a dose-dependent fashion (Fig. 3D). Importantly, the inhibitory effect was partially overcome at higher P5 concentrations [with 10 µM 19-OHT: 100% at 1 µM P5; 50% at 10 µM P5; 19.6% at 100 µM P5], consistent with competitive inhibition of HSD3B, similar to that observed with 19-Atriol [with 10 µM 19-Atriol: 100% at 1 µM P5; 47.1% at 10 µM P5; 32.2% at 100 µM P5, p<0.05] (Fig. 2C). Together, these results establish that 19-Atriol is both a substrate and a competitive inhibitor of HSD3B, and that its metabolite 19-OHT retains inhibitory activity, thereby amplifying the blockade of P4 synthesis.

### Distinct inhibitory actions of 19-Atriol and 19-OHT on upstream steps in steroidogenesis

To determine whether 19-Atriol and its metabolite 19-OHT act solely at the level of HSD3B or also influence upstream steps, we compared levels of both P4 and P5 using LC-MS/MS. Under Bt_2_cAMP stimulation, both compounds abolished P4 production in MA-10 and MA-10:*Tspo*^*Δ/Δ*^ cells (Fig. 4A). Unlike the classical HSD3B inhibitor trilostane, however, neither 19-Atriol nor 19-OHT produced an accumulation of P5 in both MA-10 and MA-10:*Tspo*^*Δ/Δ*^ cells. The absence of P5 buildup indicated that, in addition to inhibiting HSD3B, these compounds impair an upstream step required for P5 synthesis, such as CYP11A1 activity or cholesterol availability.

**Figure 4.**
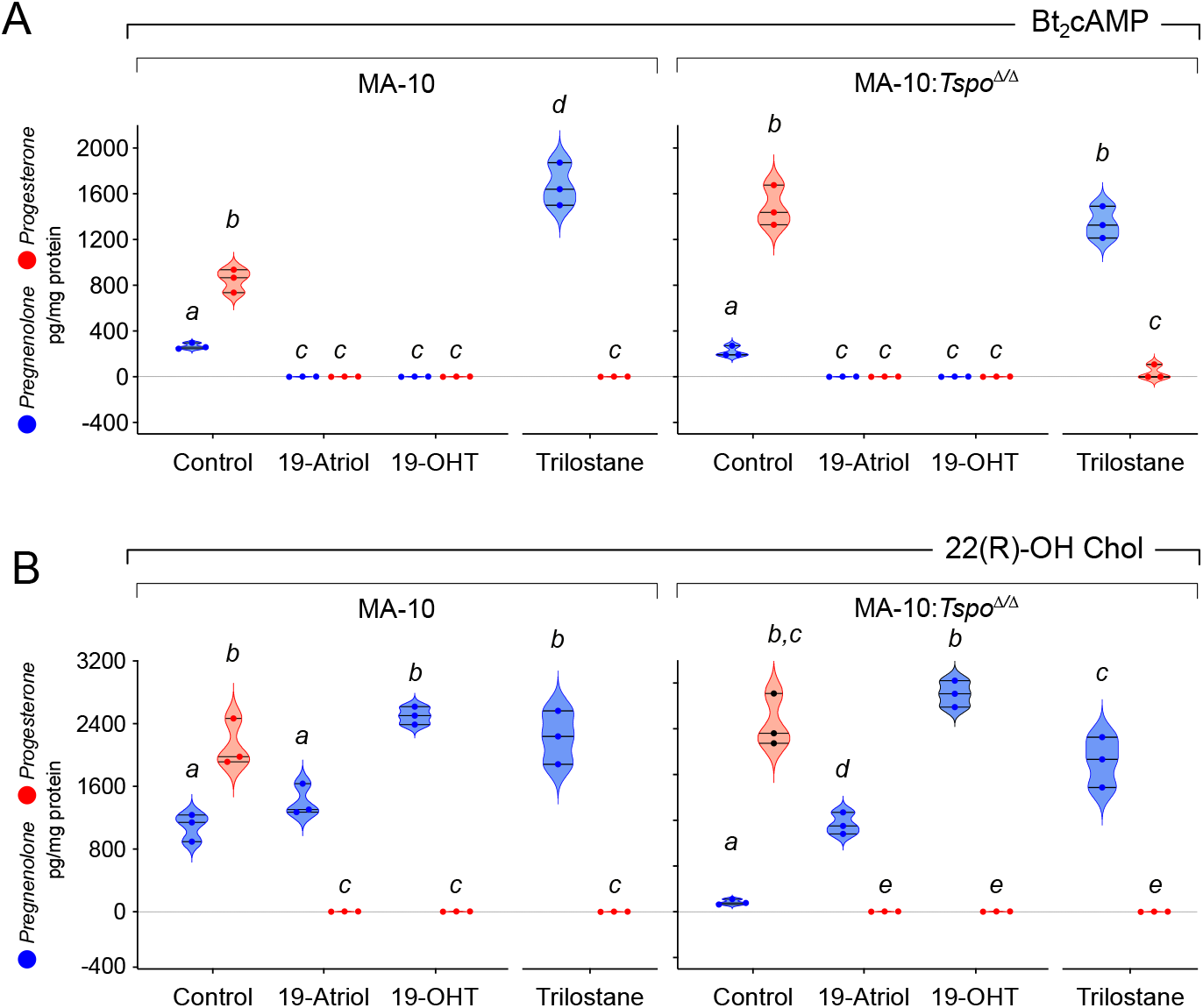
Distinct inhibitory actions of 19-Atriol and 19-OHT on HSD3B and upstream steps, independent of TSPO. **(A)** LC-MS/MS analysis revealed that under Bt_2_cAMP stimulation, both 19-Atriol and 19-OHT abolished P4 production in MA-10 and MA-10:*Tspo*^*Δ/Δ*^ cells. However, unlike the classical HSD3B inhibitor trilostane, 19-Atriol and 19-OHT did not produce a P5 buildup. The absence of P5 accumulation indicates that, in addition to inhibiting HSD3B, these treatments suppress an upstream step required for P5 formation such as CYP11A1 activity and/or cholesterol availability. **(B)** When the cholesterol-delivery step is bypassed with 22R-HC, LC-MS/MS analysis revealed that 19-Atriol and 19-OHT block HSD3B and P5 synthesis in both MA-10 and MA-10:*Tspo*^*Δ/Δ*^ cells. However, their profiles diverge: 19-Atriol causes a modest P5 accumulation, consistent with partial CYP11A1 inhibition, whereas 19-OHT produces a pronounced P5 buildup, consistent with potent HSD3B inhibition without impairing CYP11A1. Trilostane produced robust P5 accumulation with loss of P4, as expected for direct 3β-HSD blockade. The differences in these results for 19-Atriol and 19-OHT were consistent between MA-10 and MA-10:*Tspo*^*Δ/Δ*^ cells. Data are shown as violin plot distributions of replicates (points), with statistically significant differences indicated by different alphabets (p<0.05).

To further localize their actions, we bypassed cholesterol delivery by supplementing cells with the CYP11A1 intermediate 22R-HC. In this setting, both 19-Atriol and 19-OHT again suppressed P4 synthesis in wild-type and TSPO-null cells (Fig. 4B). Importantly, their inhibition profiles diverged: 19-Atriol treatment caused a modest accumulation of P5, consistent with partial inhibition of CYP11A1 activity, whereas 19-OHT treatment led to a pronounced buildup of P5, characteristic of direct HSD3B blockade without affecting CYP11A1. As expected, trilostane produced robust P5 accumulation with loss of P4, validating the assay readout. These differences between 19-Atriol and 19-OHT were reproducible across both MA-10 and MA-10:*Tspo*^*Δ/Δ*^ cells backgrounds. We also detected 20α-hydroxyprogesterone as a metabolite of P4, which followed the same inhibition pattern observed for P4 with 19-Atriol and 19-OHT (Figure S1). Collectively, these findings demonstrate that while both compounds block HSD3B, 19-Atriol additionally impinges on upstream CYP11A1 activity, whereas 19-OHT selectively inhibits HSD3B. Crucially, none of these mechanisms involved TSPO.

### CRAC motif mapping does not support a TSPO-targeting model for 19-Atriol

The original interpretation that 19-Atriol inhibits steroidogenesis through direct binding to TSPO was based on presumed engagement with a claimed cholesterol-recognition amino acid consensus (CRAC) motif at amino acids 150 to 156 of the transmembrane region 5 (TM5; indicated as CRAC-1 in Fig. 5). We have previously pointed out the loose definition of this motif (19), in that the consensus sequence (L/V–X_1–5_–Y–X_1–5_–R/K) can be detected roughly once every 112 amino acids, averaging ∼2.7 CRAC motifs per protein (41). Therefore, we mapped the motif pattern for CRAC, and an inverted version known as CARC (K/R–X_1–5_–Y–X_1–5_–L/V) to the TSPO primary sequence. From this, we detected two distinct CRAC motifs and one CARC motif distributed across the five transmembrane helices of TSPO (Fig. 5A-B). This multiplicity is consistent with statistical expectations and underscores that motif presence alone can neither predict or define a mechanistic cholesterol transport function (42). Structural modeling has previously highlighted CRAC-1 as the most likely cholesterol-binding region (43). However, the constituent side chains of this motif are distributed circumferentially along the alpha-helical turn and embedded within the polar headgroup region of the outer mitochondrial membrane (OMM) (Fig. 5B). In addition, we identify a previously unrecognized CRAC-2 motif spanning amino acids 31 to 39 within the first transmembrane helix (TM1). However, this region lies entirely outside the membrane projecting into the cytoplasm (Fig. 5B). Beyond these predictions from structure, an experimental proteome-wide cholesterol mapping study using click-chemistry identified three cholesterol-associated peptides predominantly within TM1 (residues 2–33), overlapping our *de novo* predicted CRAC-2, as well as a fourth peptide within the TM2-TM3 loop (amino acids 69 to 78) (44). More recent solid-state NMR data also demonstrate that cholesterol interactions with TSPO occur across multiple transmembrane domains in a distributed manner (45). These findings collectively argue against the existence of a discrete, high-affinity cholesterol-binding site within TSPO and specific binding of 19-Atriol to the CRAC-1 motif, which lacks structural and functional validation.

**Figure 5.**
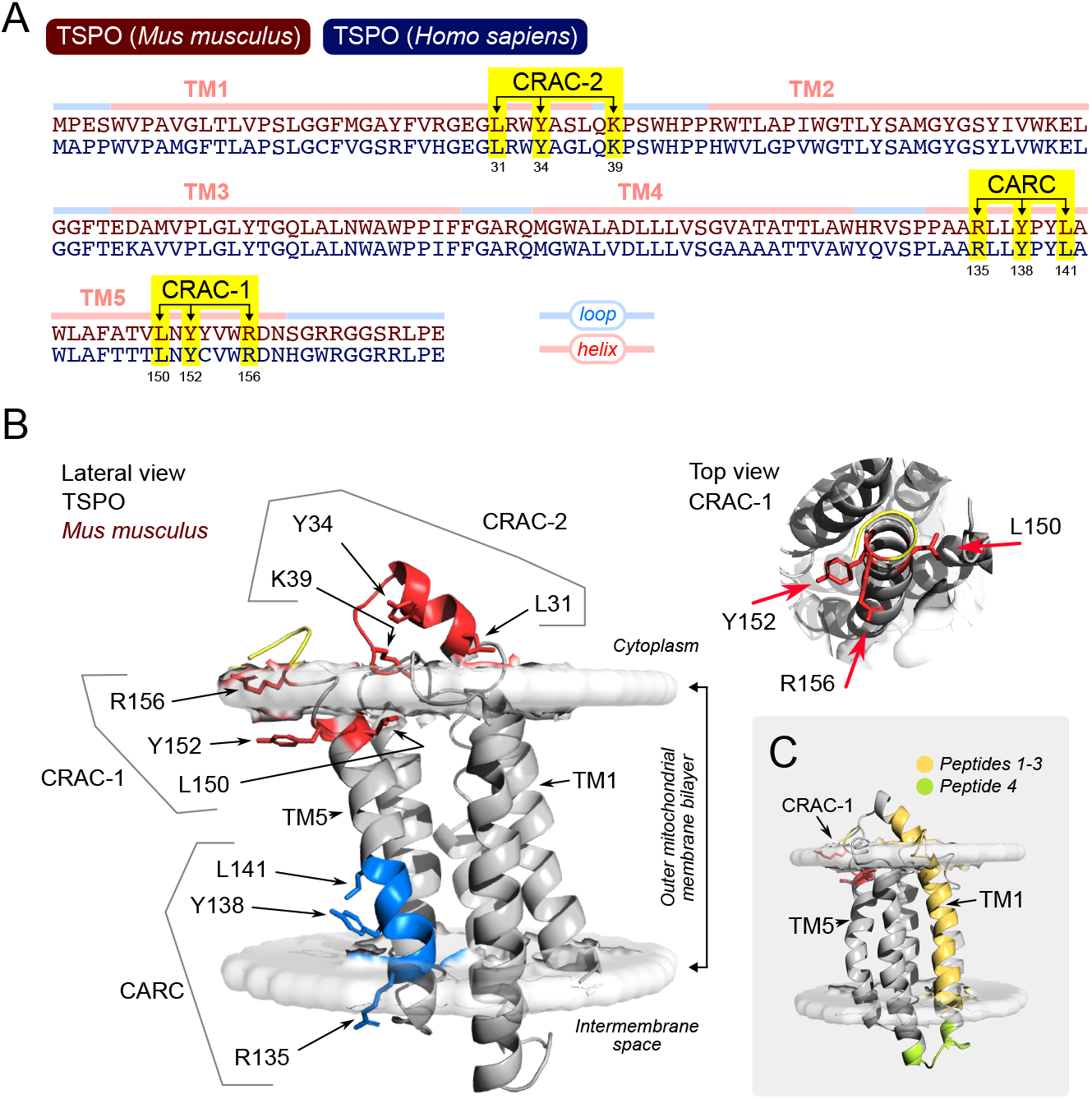
Mapping CRAC/CARC motifs and cholesterol-associating regions in TSPO. **(A)** Sequence alignment of mouse and human TSPO highlighting transmembrane helices (TM1– TM5), loops, and cholesterol-recognition motifs: CRAC-2 (aa 31–39), CRAC-1 (aa 150–156), and an inverted CARC motif (aa 135–141). **(B)** Mouse TSPO structure (PDB: 2MGY), oriented in the outer mitochondrial membrane bilayer. Left: Lateral view shows CRAC-2 projecting into the cytoplasm, while CRAC-1 and CARC are embedded within transmembrane helix TM5. Side chains for motif residues are visualized and labeled. Right: Top-down view of CRAC-1 shows circumferential side chain positioning. **(C)** Structural overlay of cholesterol-binding regions/peptides identified in a proteome-wide click-chemistry study (44). Peptides 1–3 (yellow) map to TM1, overlapping CRAC-2; peptide 4 (green) localizes to the TM2-TM3 loop. No peptides were found near CRAC-1 or CARC motifs, suggesting cholesterol association occurs outside canonical motif regions.

## Discussion

*De novo* steroid biosynthesis depends on the precise mobilization of cholesterol to the inner mitochondrial membrane (IMM), where CYP11A1 catalyzes its conversion into P5, the first steroid in the pathway (46–48). Because this conversion requires cholesterol to traverse the aqueous intermembrane space, import has long been considered the rate-limiting step of steroidogenesis (8). STAR was identified as an essential mediator of this process (21), and has now been established to function as a cholesterol shuttle within the mitochondrial intermembrane space (22). However, before the recognition of STAR’s intermembrane space localization and import-dependent mechanism, STAR was believed to act from the cytoplasm, stimulating cholesterol delivery externally to mitochondria (49). This cytoplasmic mechanism created a gap that was filled by the proposition that STAR required cooperation with TSPO, an outer membrane protein, to achieve cholesterol import (7). This model became entrenched through decades of pharmacological studies that attributed agonist or antagonist effects of TSPO ligands to changes in steroid output (4–6), albeit transient and at very low levels. The advent of *Tspo* gene-deleted cell and animal studies has since invalidated this model, showing that steroidogenesis proceeds normally in the absence of TSPO (13–17). These findings not only rule out a functional requirement for TSPO in STAR-mediated cholesterol import but also highlight major inconsistencies in how TSPO’s role in steroidogenesis was defined (19, 20). With STAR’s mechanism now clarified and TSPO excluded as a cholesterol transport factor, defined genetic models offer a rigorous framework for reassessing the pharmacology of TSPO-binding compounds currently under consideration for diagnostic and therapeutic use (50, 51).

The attribution of pharmacological activity from TSPO-binding compounds to steroid hormone production has persisted, despite two important considerations, which these interpretations neglect. First, even in steroidogenic tissues such as the adrenal cortex and gonads, putative TSPO agonists such as PK11195 elicit only modest and transient increases in steroid output typically ∼40-fold lower (2.5 % level) than that induced by signal transducers such as Bt_2_cAMP (15). Second, across both adrenal and neuronal systems, the prototypical TSPO-binding compound, PK11195 has frequently produced antagonistic rather than agonistic effects on steroidogenesis (52, 53). These inconsistencies have made it difficult to reconcile the physiological significance of PK11195 responses, particularly in light of its known actions on other cellular targets, including the constitutive androstane receptor (54) and the F_1_F_0_ ATP synthase (55). Our own studies using CRISPR/Cas9-mediated *Tspo*-deleted MA-10 cells demonstrated that the transient effects of PK11195 on modest steroidogenic stimulation are fully retained in the absence of TSPO (15), directly refuting the hypothesis that TSPO mediates this pharmacology. Collectively, these findings correct decades of TSPO research (56–58), and underscore the necessity of genetic validation when evaluating the specificity of TSPO-targeting drugs (19).

The original report of 19-Atriol effects on steroidogenesis (37) attributed its inhibition of P4 synthesis in MA-10 cells to direct binding at a CRAC motif of TSPO (59, 60), thereby blocking cholesterol import into mitochondria. While we fully reproduce the observed inhibition of P4 synthesis, our findings decisively demonstrate that this effect is entirely independent of TSPO, as identical inhibition profiles were obtained in TSPO null MA-10:*Tspo*^*Δ/Δ*^ cells. This disconnect underscores the broader conceptual problem of linking cholesterol-associating motifs on TSPO to cholesterol transport function and *de novo* steroidogenesis. The CRAC motif, first described on TSPO (43, 60), is a statistically common pattern across the proteome, and its presence does not equate cholesterol association (41, 42). Indeed, our prediction of CRAC motifs reveals not one, but two CRAC motifs and one inverted CARC motif in TSPO. Although CRAC-1 in TM5 has been posited as the putative cholesterol binding site (60), proteome-wide cholesterol interactome profiling using click-chemistry instead identified cholesterol-associated peptides overlapping the TM1 region near our predicted CRAC-2 (44). Structural modeling also shows that CRAC/CARC side chains are consistent with nonspecific cholesterol association in lipid-raft-like domains (61, 62). More recent NMR studies have also demonstrated cholesterol association on TSPO is not confined to a single motif such as CRAC-1 (45). Collectively, these data reveal that the assignment of functional significance to CRAC and CARC motifs within proteins, particularly to CRAC-1 of TSPO, is not reliable (42). Importantly, our results show that any putative interaction between 19-Atriol and TSPO, across CRAC or CARC motifs, is irrelevant to its mechanism of steroidogenic inhibition. Instead, 19-Atriol and its active metabolite 19-OHT act through direct competition with steroidogenic enzymes, primarily HSD3B. Thus, prior claims linking its activity to binding CRAC-1 of TSPO and mitochondrial cholesterol transport reflect a mechanistic misattribution from a flawed assumption.

The structural properties of 19-Atriol and its metabolite 19-OHT provide a credible rationale for their inhibitory actions across multiple steps of the steroidogenic pathway. 19-Atriol retains the Δ^5^ double bond and 3β-hydroxyl group characteristic of P5, while introducing a 19-hydroxyl substitution that both preserves sterol-like recognition and introduces steric and polarity changes within the active site of enzymes that use cholesterol and P5 substrates. These features allow 19-Atriol to act as a competitive substrate for HSD3B, directly inhibiting the occupancy of P5, and through similar occupancy, it also partially impedes CYP11A1-mediated side-chain cleavage of cholesterol. The formation of 19-OHT by HSD3B further amplifies these effects, as HSD3B enzymes are known to show product inhibition (63). This 19-OHT metabolite structurally resembles testosterone but with an additional hydroxyl at C19, enabling it to bind the HSD3B active site and inhibit enzymatic turnover without significantly engaging CYP11A1. The differential response we observed with 22R-HC supplementation underscores this interpretation: because 22R-HC has much higher affinity for CYP11A1 than cholesterol (64), its utilization is less susceptible to inhibition, revealing that 19-Atriol interferes with cholesterol access or orientation rather than strongly competing at the catalytic site itself. This divergence explains why 19-Atriol produces modest P5 accumulation (indicative of partial CYP11A1 inhibition) whereas 19-OHT causes pronounced P5 buildup consistent with selective HSD3B blockade. Beyond enzyme inhibition, the sterol-like backbone of 19-Atriol could also compete with endogenous cholesterol for binding pockets involved in mitochondrial delivery of cholesterol, providing an additional mechanism for reduced P5 synthesis. Together, these structural considerations explain the distinct but complementary inhibitory profiles of 19-Atriol and 19-OHT across cholesterol import, side-chain cleavage, and P5 metabolism observed in our study. These mechanistic insights, in turn, provide a framework to reinterpret earlier reports of 19-Atriol pharmacology in physiological and disease models.

In the ocular setting, *ex vivo* experiments in rat retina demonstrated that allopregnanolone biosynthesis mitigates the damaging effects of elevated intraocular pressure (65). In this context, 19-Atriol was shown to suppress allopregnanolone production and exacerbate pressure-induced retinal injury, outcomes that were originally ascribed to TSPO antagonism. Based on our results, these effects can instead be explained by direct inhibition of steroidogenic enzymes, thereby reducing neurosteroid synthesis. This reinterpretation not only accounts for the previously observed retinal toxicity but also emphasizes that neurosteroid availability rather than TSPO expression is the critical determinant of neuronal resilience in glaucoma models.

Looking forward, these findings underscore the need to reassess decades of TSPO pharmacology using defined genetic models and rigorous biochemical validation. The implications extend beyond reinterpretation of prior work on steroidogenesis in ocular and neuronal disease models, to the ongoing evaluation of TSPO-binding drugs that continue to advance toward clinical application. Future efforts should integrate structural, genetic, and functional approaches to establish authentic target-mechanism relationships, enabling pharmacological discovery to progress with greater precision and translational impact.

## Experimental procedures

### Synthesis of 19-Atriol

19-Acetoxy-androst-5-en-3β,17β-diol was synthesized from 19-hydroxy-androst-4-ene-3,17-dione following a two-step sequence. To a solution of 19-hydroxy-androst-4-ene-3,17-dione (0.5 g, 1.67 mmol) in acetic anhydride (6 ml) was added sodium iodide (7 mmol) and chlorotrimethylsilane (7 mmol) at 0°C under N_2_. The reaction was poured into aqueous NaHCO_3_ and the 3,19-(diacetoxy)-androsta-3,5-dien-17-one product was extracted into EtOAc (300 ml). The extract was washed with brine (100 mL × 2), dried over anhydrous Na_2_SO_4_, filtered, the solvent removed, and the residue was dissolved in EtOH (50 ml). NaBH_4_ was added and the reaction was stirred at 23°C for 16 h. Aqueous NH_4_Cl was added and stirred for 10 min. Most of EtOH was removed under reduced pressure and the product was extracted into EtOAc (150 ml × 2) from the remaining aqueous solution. The combined extracts were dried over anhydrous Na_2_SO_4_, filtered, the solvent removed, and the residue was recrystallized from Et_2_O/CH_2_Cl_2_/hexanes (v/v/v=10/2/10) to afford the product 19-Acetoxy-androst-5-en-3β,17β-diol (19-OHT; 135 mg, 26%). Structure was confirmed by ^1^H and ^13^C NMR spectra: ^1^H NMR (400 MHz, CDCl3) δ 5.56-5.55 (m, 1H), 4.45 (d, J = 12.1 Hz, 1H), 3.95 (d, J = 12.1 Hz, 1H), 3.60-3.41 (m, 2H), 2.32-0.83 (m, 21H), 2.01 (s, 3H), 0.73 (s, 3H); 13C NMR (100 MHz, CDCl3) δ 170.9, 135.7, 125.3, 81.6, 71.2, 64.7, 51.8, 50.2, 42.8, 42.2, 39.6, 36.7, 33.8, 32.9, 31.6, 30.9, 30.3, 23.3, 21.2, 21.1, 11.1.

19-Atriol was prepared from 19-acetoxy-androst-5-en-3β,17β-diol. A solution of 19-acetoxy-androst-5-en-3β,17β-diol (130 mg, 0.37 mmol) in MeOH (15 ml) was treated with K_2_CO_3_ (552 mg, 4 mmol) at 23°C. The reaction mixture was refluxed for 16 h, cooled, and concentrated under reduced pressure. The residue was purified by flash column chromatography on silica gel (silica gel, eluted with 10% MeOH in CH_2_Cl_2_) to give Androst-5-ene-3β,17β,19-triol (19-Atriol; 87 mg, 77%). Structure was confirmed by ^1^H and ^13^C NMR spectra: ^1^H NMR (400 MHz, CDCl_3_-CD_3_OD) δ 5.22-5.21 (m, 1H), 3.94 (d, J = 11.4, 1H), 3.70-3.46 (m, 3H), 3.42 (s, 1H), 2.44-0.93 (m, 21H), 0.90 (s, 3H); ^13^C NMR (100 MHz, CDCl_3_-CD_3_OD) δ 135.9,124.5, 80.6, 70.3, 61.6, 51.5, 50.2, 42.1, 41.2, 40.6, 36.3, 32.7, 32.4, 30.6, 30.3, 28.8, 22.4, 20.7, 10.1.

### MA-10 Cell culture and treatments

TSPO-deleted MA-10 clones (MA-10:*Tspo*^*Δ/Δ*^) were previously generated and validated (15). MA-10 Leydig cells and MA-10:*Tspo*^*Δ/Δ*^ cells were cultured in DMEM supplemented with 10% fetal bovine serum, 1% penicillin-streptomycin, and 1% non-essential amino acids following established protocols (66). For experimental treatments, cells were plated in 96-well plates pre-coated with 0.1% gelatin at a density of 5 × 10^4^ cells per well and allowed to attach overnight. Cells were stimulated with either 0.5 mM Bt_2_cAMP (Sigma) or 20 µM 22(R)-hydroxycholesterol (22R-HC; Sigma) for 3 h to induce steroidogenesis.

To determine steroidogenic steps inhibited by 19-Atriol, MA-10 and MA-10:*Tspo*^*Δ/Δ*^ cells were treated with 1, 10, or 100 µM 19-Atriol under Bt_2_cAMP stimulation or 22R-HC treatment. Parallel experiments were performed with the 19-Atriol metabolite, 19-hydroxytestosterone (19-OHT), at identical concentrations. After treatments, supernatants were collected to quantify steroid hormones, and cells were harvested for protein estimation. Each set of treatments was performed in duplicate 96-well plates, yielding 200 µl supernatant per condition. Of this, 150 µl was analyzed by mass spectrometry, while 50 µl was assayed for P5 and P4 levels using radioimmunoassay (RIA) or LC-MS/MS as described. In parallel, cells were harvested for protein quantification using the bicinchoninic acid (BCA) method (Pierce™ BCA Protein Assay Kit, Thermo Fisher Scientific) for normalization.

### 3β-HSD activity assay

To establish an *in vitro* assay for 3β-hydroxysteroid dehydrogenase (HSD3B) activity, MA-10 and MA-10:*Tspo*^*Δ/Δ*^ cells were supplemented with P5 (1, 5, 10, or 100 µM) for 3 h to monitor its conversion to P4. This approach bypasses the CYP11A1 step and directly assays HSD3B function. Following incubation, culture supernatants were collected and stored at -20°C for subsequent P4 quantification by RIA. For inhibition testing, cells we co-treated with 19-Atriol or 19-OHT (10 or 100 µM) during the incubation period. In parallel, cells were harvested for protein quantification using the BCA method. As a pharmacologic control, trilostane (20 µM), a well-characterized HSD3B inhibitor, was included under identical assay conditions.

### Radioimmunoassay for progesterone

Progesterone levels in culture supernatants were quantified as previously described (15, 66). Briefly, supernatants were incubated overnight at 4 °C with ^125^I-labeled progesterone (MP Biomedicals) and an anti-progesterone antibody (67) for competitive binding. The free fraction was removed by adding a charcoal–dextran suspension, followed by a 10 min incubation at 4 °C and centrifugation at 1700 g for 10 min. Radioactivity in the bound fraction was measured using a scintillation counter (Clinigamma Automatic, Wallac). Progesterone concentrations were determined from standard curves generated with serial progesterone standards and normalized to total protein content per well.

### Steroid quantitation by Liquid Chromatography Tandem Mass-Spectrometry (LC-MS/MS)

Samples were prepared by combining 0.1 ml media, 0.1 ml deionized water, and 0.1 ml internal standards. The mixtures were added into supported liquid extraction columns (Biotage Isolute, 820-0055-B) and absorbed for 5 min. Steroids were eluted with 1.8 ml methyl-tert-butyl ether, dried with a vacuum evaporator centrifuge, and reconstituted in 0.2 ml of 40:60 methanol:water (v:v). Aliquots (10 μl) were injected via autosampler and resolved using two-dimensional liquid chromatography, starting with on-line cleanup using a Hypersil GOLD C4 column (Thermo, 3 × 10 mm, 25503-01300) and an Agilent 1260 HPLC in the first dimension. The eluting steroids were routed onto a Kinetex biphenyl column (Phenomenex 2.1 × 50 mm, 00B-4622-AN) using a 10-port switching valve and resolved using an Agilent 1290 UPLC with water and methanol gradients containing 0.2 mmol/L ammonium fluoride (Sigma, CAS 12125-01-8) for the second dimension. The second column effluent was directed into the source of an Agilent 6495A triple quadrupole tandem mass spectrometer for electrospray ionization, and steroids were quantified with dynamic multiple reaction monitoring (MRM) to detect analytes and internal standards (see Supporting Information Tables S1-S3) as described (68).

### Immunoblots

Cells were lysed in RIPA buffer supplemented with protease inhibitors (Sigma). Equal amounts of protein lysates were resolved by SDS–PAGE and transferred to PVDF membranes. Membranes were probed with rabbit monoclonal antibodies against TSPO (Abcam) and CYP11A1 (Cell Signaling Technology), and a rabbit polyclonal antibody against STAR (66). A monoclonal mouse antibody against β-actin (LI-COR) was used as a loading control. Fluorescent detection of TSPO, CYP11A1, STAR, and β-actin was carried out using species-specific IRDye 700 and IRDye 800 secondary antibodies (LI-COR), and visualized on a quantitative imaging system (C600, Azure Biosystems).

### TSPO structure

Structural features of the murine TSPO protein were analyzed using a combination of experimental and computational approaches. The murine TSPO NMR structure (PDB: 2MGY) was used (69). Mitochondrial outer membrane orientation was from the OPM database (70). All structural visualization and annotations were performed in PyMOL 2.6 (pymol.org).

### Statistics

All experiments were performed with a minimum of three independent replicates, and each experiment was reproduced at least twice to ensure technical rigor. Quantitative comparisons between two groups were assessed using Student’s t-test. For comparisons involving more than two groups, one-way ANOVA followed by Tukey’s post hoc test was applied. Statistical analyses were performed in GraphPad Prism 5, and differences were considered significant at p<0.05.

## Acknowledgements

We thank Dr. Mario Ascoli (University of Iowa) for providing the MA-10 Leydig cell line and Mr. David Stouffer for assistance with mass spectrometry methods. This work was supported by the National Institutes of Health grant R01 DK110059 to V.S., R01 GM086596 to R.J.A., and R01 MH122379 to D.C.

## Supplementary Information

**Figure S1.**
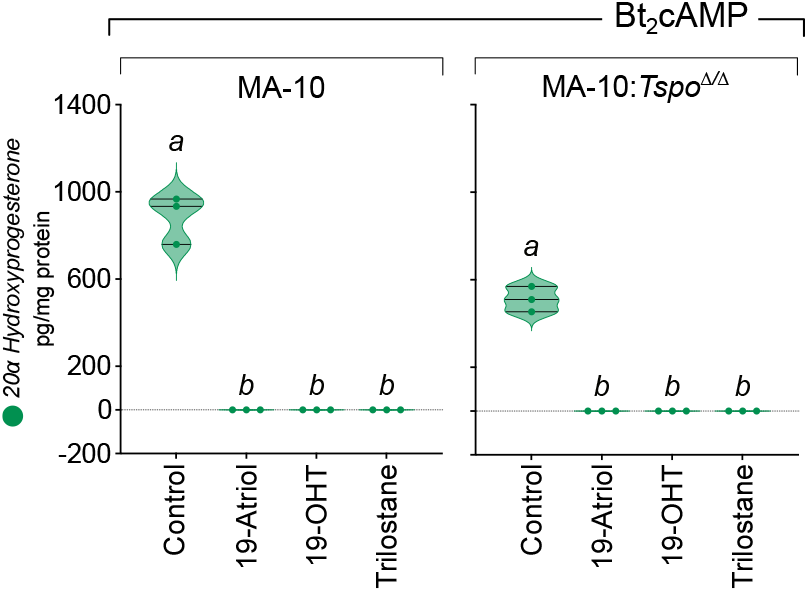
Detection of 20α-hydroxyprogesterone (20α-OHP) as a metabolite of P4 in MA-10 cells. LC–MS/MS analysis also identified 20α-OHP (same precursor and product ions as pregnenolone) in culture supernatants of MA-10 cells. The 20α-OHP is formed from P4 by 20α-hydroxysteroid dehydrogenase (20α-HSD; AKR1C family), and its abundance paralleled the inhibition patterns of P4 observed with 19-Atriol and 19-OHT treatment, consistent with blockade of HSD3B activity. Data are shown as violin plot distributions of replicates (points), with statistically significant differences indicated by different alphabets (p<0.05).

**Table S1.** Liquid chromatography parameters used for binary pump tuning. Column Temperature was maintained at 30°C. The Switch valve was moved from position 1 to position 2 between 0.50 and 03.20 minutes. Mobile Phase A: Water, Mobile Phase B: Methanol; both contain 0.20 mM Ammonium Fluoride

| Binary Pump 1 | Step | Time (min) | Water A% | Methanol B% | Flow (mL/min) |
| --- | --- | --- | --- | --- | --- |
|  | 1 | 0.7 | 65 | 35 | 0.5 |
|  | 2 | 0.71 | 0 | 100 | 0.25 |
|  | 3 | 3.4 | 0 | 100 | 0.25 |
|  | 4 | 3.41 | 0 | 100 | 0.5 |
|  | 5 | 4.4 | 0 | 100 | 0.5 |
|  | 6 | 4.41 | 65 | 35 | 0.5 |

| Binary Pump 2 | Step | Time (min) | Water A% | Methanol B% | Flow (mL/min) |
| --- | --- | --- | --- | --- | --- |
|  | 1 | 0.5 | 68 | 32 | 0.25 |
|  | 2 | 0.51 | 68 | 32 | 0.25 |
|  | 3 | 1.6 | 55 | 45 | 0.25 |
|  | 4 | 1.61 | 45 | 55 | 0.25 |
|  | 5 | 8 | 5 | 95 | 0.25 |
|  | 6 | 8.5 | 0 | 100 | 0.5 |
|  | 7 | 9 | 0 | 100 | 0.5 |
|  | 8 | 9.01 | 68 | 32 | 0.5 |

**Table S2.**
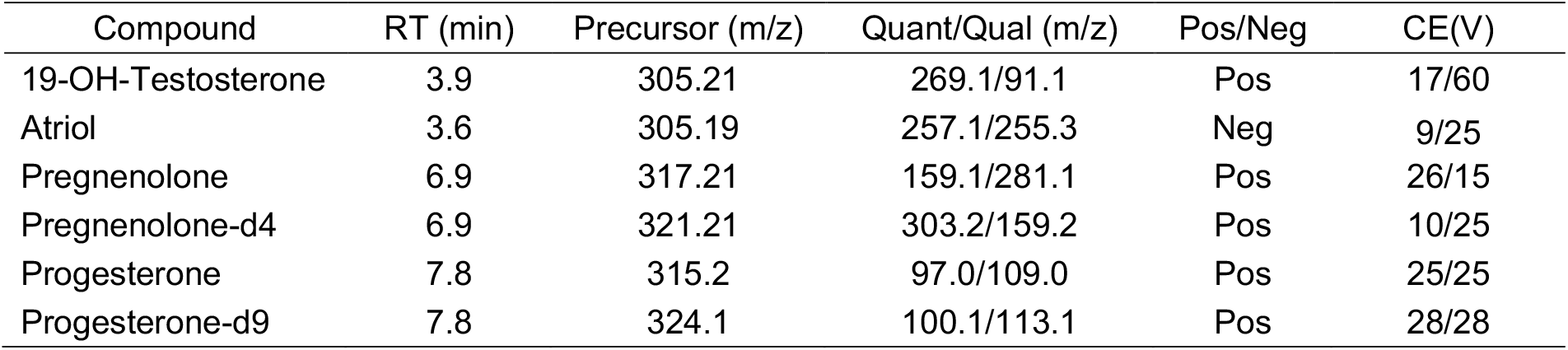
Mass spectrometry (MS/MS) parameters used for targeted steroid hormone quantification. Retention times (RT), precursor and product ion transitions (quantifier/qualifier), ionization polarity, and collision energies (CE) are shown for each compound

**Table S3.** Source parameters used for electron spray ionization.

| Parameter | Value |
| --- | --- |
| Gas Temperature | 120° C |
| Gas Flow | 16 L/min |
| Nebulizer | 45 psi |
| Sheath Gas Temperature | 200° C |
| Sheath Gas Flow | 12 L/min |
| Capillary Voltage | +2650 V |
| Nozzle Voltage | +1000 V |
| Ion Funnel (high) | 150/-100 |
| Ion Funnel (low) | 120/-70 |

## References

1. Gavish, M., Bachman, I., Shoukrun, R., Katz, Y., Veenman, L., Weisinger, G., and Weizman, A. (1999) Enigma of the peripheral benzodiazepine receptor. Pharmacological Reviews. 51, 629–650

2. Selvaraj, V., and Stocco, D. M. (2015) The changing landscape in translocator protein (TSPO) function. Trends in Endocrinology and Metabolism. 26, 341–348

3. Anholt, R. R. H., Pedersen, P. L., De Souza, E. B., and Snyder, S. H. (1986) The Peripheral-type Benzodiazepine Receptor, Localization to the mitochondrial outer membrane. Journal of Biological Chemistry. 261, 576–583

4. Mukhin, A. G., Papadopoulos, V., Costa, E., and Krueger, K. E. (1989) Mitochondrial benzodiazepine receptors regulate steroid biosynthesis. Proceedings of the National Academy of Sciences of the United States of America. 86, 9813–9816

5. Papadopoulos, V., Mukhin, A. G., Costa, E., and Krueger, K. E. (1990) The peripheral-type benzodiazepine receptor is functionally linked to Leydig cell steroidogenesis. J Biol Chem. 265, 3772–3779

6. Krueger, K. E., and Papadopoulos, V. (1990) Peripheral-type benzodiazepine receptors mediate translocation of cholesterol from outer to inner mitochondrial membranes in adrenocortical cells. The Journal of Biological Chemistry. 265, 15015–15022

7. Hauet, T., Yao, Z. X., Bose, H. S., Wall, C. T., Han, Z., Li, W., Hales, D. B., Miller, W. L., Culty, M., and Papadopoulos, V. (2005) Peripheral-type benzodiazepine receptor-mediated action of steroidogenic acute regulatory protein on cholesterol entry into leydig cell mitochondria. Molecular Endocrinology. 19, 540–554

8. Stocco, D. M., Zhao, A. H., Tu, L. N., Morohaku, K., and Selvaraj, V. (2017) A brief history of the search for the protein(s) involved in the acute regulation of steroidogenesis. Molecular and Cellular Endocrinology. 441, 7–16

9. Papadopoulos, V., and Lecanu, L. (2009) Translocator protein (18 kDa) TSPO: an emerging therapeutic target in neurotrauma. Experimental neurology. 219, 53–57

10. Rupprecht, R., Papadopoulos, V., Rammes, G., Baghai, T. C., Fan, J., Akula, N., Groyer, G., Adams, D., and Schumacher, M. (2010) Translocator protein (18 kDa) (TSPO) as a therapeutic target for neurological and psychiatric disorders. Nature Reviews Drug Discovery. 9, 971–988

11. Qi, X., Xu, J., Wang, F., and Xiao, J. (2012) Translocator protein (18 kDa): a promising therapeutic target and diagnostic tool for cardiovascular diseases. Oxidative medicine and cellular longevity. 2012, 162934

12. Vlodavsky, E., Palzur, E., and Soustiel, J. F. (2013) 18 kDa Translocator Protein as a Potential Therapeutic Target for Traumatic Brain Injury. CNS & neurological disorders drug targets

13. Morohaku, K., Pelton, S. H., Daugherty, D. J., Butler, W. R., Deng, W., and Selvaraj, V. (2014) Translocator Protein/Peripheral Benzodiazepine Receptor Is Not Required for Steroid Hormone Biosynthesis. Endocrinology. 155, 89–97

14. Tu, L. N., Morohaku, K., Manna, P. R., Pelton, S. H., Butler, W. R., Stocco, D. M., and Selvaraj, V. (2014) Peripheral benzodiazepine receptor/translocator protein global knock-out mice are viable with no effects on steroid hormone biosynthesis. Journal of Biological Chemistry. 289, 27444–27454

15. Tu, L. N., Zhao, A. H., Stocco, D. M., and Selvaraj, V. (2015) PK11195 effect on steroidogenesis is not mediated through the translocator protein (TSPO). Endocrinology. 156, 1033–1039

16. Banati, R. B., Middleton, R. J., Chan, R., Hatty, C. R., Wai-Ying Kam, W., Quin, C., Graeber, M. B., Parmar, A., Zahra, D., Callaghan, P., Fok, S., Howell, N. R., Gregoire, M., Szabo, A., Pham, T., Davis, E., and Liu, G. J. (2014) Positron emission tomography and functional characterization of a complete PBR/TSPO knockout. Nature Communications. 10.1038/ncomms6452

17. Wang, H., Zhai, K., Xue, Y., Yang, J., Yang, Q., Fu, Y., Hu, Y., Liu, F., Wang, W., Cui, L., Chen, H., Zhang, J., and He, W. (2016) Global Deletion of TSPO Does Not Affect the Viability and Gene Expression Profile. PLOS ONE. 11, e0167307

18. Daugherty, D. J., Chechneva, O., Mayrhofer, F., and Deng, W. (2016) The hGFAP-driven conditional TSPO knockout is protective in a mouse model of multiple sclerosis. Scientific Reports. 10.1038/srep22556

19. Selvaraj, V., Stocco, D. M., and Tu, L. N. (2015) Minireview: translocator protein (TSPO) and steroidogenesis: a reappraisal. Molecular Endocrinology. 29, 490–501

20. Selvaraj, V., and Tu, L. N. (2016) Current status and future perspectives: TSPO in steroid neuroendocrinology. J Endocrinol. 231, R1–R30

21. Clark, B. J., Wells, J., King, S. R., and Stocco, D. M. (1994) The purification, cloning, and expression of a novel luteinizing hormone-induced mitochondrial protein in MA-10 mouse Leydig tumor cells. Characterization of the steroidogenic acute regulatory protein (StAR). The Journal of biological chemistry. 269, 28314–28322

22. Koganti, P. P., Zhao, A. H., Kern, M. C., Fassinou, A. C. R., and Selvaraj, V. (2025) STAR/STARD1: a mitochondrial intermembrane space cholesterol shuttle degraded through mitophagy. bioRxiv. 10.1101/2025.04.03.647084

23. Smith, C. R., Tu, L. N., Koganti, P., and Selvaraj, V. (2025) TSPO deficiency reveals a novel role for porphyrins in regulating white adipose tissue lipid metabolism. 10.1101/2025.09.14.676078

24. Tu, L. N., Zhao, A. H., Hussein, M., Stocco, D. M., and Selvaraj, V. (2016) Translocator Protein (TSPO) Affects Mitochondrial Fatty Acid Oxidation in Steroidogenic Cells. Endocrinology. 157, 1110–1121

25. Thai, P., Herren, A. W., Ren, L., Bers, D. M., Schaefer, S., and Dedkova, E. N. (2020) Mitochondrial Translocator Protein (TSPO) Prevents Heart Failure by Increasing Cardiac Utilization of Fatty Acids. Biophysical Journal. 118, 445a–446a

26. Morohaku, K., Phuong, N. S., and Selvaraj, V. (2013) Developmental Expression of Translocator Protein/Peripheral Benzodiazepine Receptor in Reproductive Tissues. PLoS ONE. 10.1371/journal.pone.0074509

27. Koganti, P. P., and Selvaraj, V. (2020) Lack of adrenal TSPO/PBR expression in hamsters reinforces correlation to triglyceride metabolism. Journal of Endocrinology. 10.1530/JOE-20-0189

28. Gudasheva, T. A., Deeva, O. A., Pantileev, A. S., Mokrov, G. V., Rybina, I. V., Yarkova, M. A., and Seredenin, S. B. (2020) The New Dipeptide TSPO Ligands: Design, Synthesis and Structure-Anxiolytic Activity Relationship. Molecules. 25, 5132

29. Corsi, F., Baglini, E., Barresi, E., Salerno, S., Cerri, C., Martini, C., Da Settimo Passetti, F., Taliani, S., Gargini, C., and Piano, I. (2022) Targeting TSPO Reduces Inflammation and Apoptosis in an In Vitro Photoreceptor-Like Model of Retinal Degeneration. ACS Chem Neurosci. 13, 3188–3197

30. Zeno, S., Zaaroor, M., Leschiner, S., Veenman, L., and Gavish, M. (2009) CoCl2 Induces Apoptosis via the 18 kDa Translocator Protein in U118MG Human Glioblastoma Cells. Biochemistry. 48, 4652–4661

31. Yasin, N., Veenman, L., Singh, S., Azrad, M., Bode, J., Vainshtein, A., Caballero, B., Marek, I., and Gavish, M. (2017) Classical and Novel TSPO Ligands for the Mitochondrial TSPO Can Modulate Nuclear Gene Expression: Implications for Mitochondrial Retrograde Signaling. Int J Mol Sci. 18, 786

32. Gatliff, J., East, D. A., Singh, A., Alvarez, M. S., Frison, M., Matic, I., Ferraina, C., Sampson, N., Turkheimer, F., and Campanella, M. (2017) A role for TSPO in mitochondrial Ca2+ homeostasis and redox stress signaling. Cell Death Dis. 8, e2896

33. Liu, G. J., Middleton, R. J., Kam, W. W. Y., Chin, D. Y., Hatty, C. R., Chan, R. H. Y., and Banati, R. B. (2017) Functional gains in energy and cell metabolism after TSPO gene insertion. Cell Cycle. 16, 436–447

34. Loth, M. K., Guariglia, S. R., Re, D. B., Perez, J., de Paiva, V. N., Dziedzic, J. L., Chambers, J. W., Azzam, D. J., and Guilarte, T. R. (2020) A Novel Interaction of Translocator Protein 18 kDa (TSPO) with NADPH Oxidase in Microglia. Mol Neurobiol. 57, 4467–4487

35. Gatliff, J., East, D., Crosby, J., Abeti, R., Harvey, R., Craigen, W., Parker, P., and Campanella, M. (2014) TSPO interacts with VDAC1 and triggers a ROS-mediated inhibition of mitochondrial quality control. Autophagy. 10, 2279–2296

36. Choi, J., Ifuku, M., Noda, M., and Guilarte, T. R. (2011) Translocator protein (18 kDa)/peripheral benzodiazepine receptor specific ligands induce microglia functions consistent with an activated state. Glia. 59, 219–230

37. Midzak, A., Akula, N., Lecanu, L., and Papadopoulos, V. (2011) Novel androstenetriol interacts with the mitochondrial translocator protein and controls steroidogenesis. J Biol Chem. 286, 9875–9887

38. Midzak, A., Rammouz, G., and Papadopoulos, V. (2012) Structure–activity relationship (SAR) analysis of a family of steroids acutely controlling steroidogenesis. Steroids. 77, 1327–1334

39. Chung, J. Y., Chen, H., Midzak, A., Burnett, A. L., Papadopoulos, V., and Zirkin, B. R. (2013) Drug ligand-induced activation of translocator protein (TSPO) stimulates steroid production by aged brown Norway rat Leydig cells. Endocrinology. 154, 2156–2165

40. Potts, G. O., Creange, J. E., Harding, H. R., and Schane, H. P. (1978) Trilostane, an orally active inhibitor of steroid biosynthesis. Steroids. 32, 257–267

41. Palmer, M. (2004) Cholesterol and the activity of bacterial toxins. FEMS microbiology letters. 238, 281–289

42. Jaipuria, G., Ukmar-Godec, T., and Zweckstetter, M. (2018) Challenges and approaches to understand cholesterol-binding impact on membrane protein function: an NMR view. Cell. Mol. Life Sci. 75, 2137–2151

43. Li, H., and Papadopoulos, V. (1998) Peripheral-type benzodiazepine receptor function in cholesterol transport. Identification of a putative cholesterol recognition/interaction amino acid sequence and consensus pattern. Endocrinology. 139, 4991–4997

44. Hulce, J. J., Cognetta, A. B., Niphakis, M. J., Tully, S. E., and Cravatt, B. F. (2013) Proteome-wide mapping of cholesterol-interacting proteins in mammalian cells. Nat Methods. 10, 259–264

45. Jaipuria, G., Giller, K., Leonov, A., Becker, S., and Zweckstetter, M. (2018) Insights into Cholesterol/Membrane Protein Interactions Using Paramagnetic Solid-State NMR. Chemistry – A European Journal. 24, 17606–17611

46. Greengard, P., Psychoyos, S., Tallan, H. H., Cooper, D. Y., Rosenthal, O., and Estabrook, R. W. (1967) Aldosterone synthesis by adrenal mitochondria. III. Participation of cytochrome P-450. Archives of Biochemistry and Biophysics. 121, 298–303

47. Omura, T., Sanders, E., Estabrook, R. W., Cooper, D. Y., and Rosenthal, O. (1966) Isolation from adrenal cortex of a nonheme iron protein and a flavoprotein functional as a reduced triphosphopyridine nucleotide-cytochrome P-450 reductase. Archives of Biochemistry and Biophysics. 117, 660–673

48. Selvaraj, V., Stocco, D. M., and Clark, B. J. (2018) Current knowledge on the acute regulation of steroidogenesis. Biol Reprod. 99, 13–26

49. Bose, H. S., Lingappa, V. R., and Miller, W. L. (2002) Rapid regulation of steroidogenesis by mitochondrial protein import. Nature. 417, 87–91

50. Kim, T. H., and Pae, A. N. (2016) Translocator protein (TSPO) ligands for the diagnosis or treatment of neurodegenerative diseases: a patent review (2010–2015; part 2). Expert Opinion on Therapeutic Patents. 26, 1353–1366

51. Kim, T. H., and Pae, A. N. (2016) Translocator protein (TSPO) ligands for the diagnosis or treatment of neurodegenerative diseases: a patent review (2010–2015; part 1). Expert Opinion on Therapeutic Patents. 26, 1325–1351

52. Romeo, E., Cavallaro, S., Korneyev, A., Kozikowski, A. P., Ma, D., Polo, A., Costa, E., and Guidotti, A. (1993) Stimulation of brain steroidogenesis by 2-aryl-indole-3-acetamide derivatives acting at the mitochondrial diazepam-binding inhibitor receptor complex. The Journal of pharmacology and experimental therapeutics. 267, 462–471

53. Cavallaro, S., Korneyev, A., Guidotti, A., and Costa, E. (1992) Diazepam-binding inhibitor (DBI)-processing products, acting at the mitochondrial DBI receptor, mediate adrenocorticotropic hormone-induced steroidogenesis in rat adrenal gland. Proceedings of the National Academy of Sciences of the United States of America. 89, 10598–10602

54. Li, L., Chen, T., Stanton, J. D., Sueyoshi, T., Negishi, M., and Wang, H. (2008) The peripheral benzodiazepine receptor ligand 1-(2-chlorophenyl-methylpropyl)-3-isoquinoline-carboxamide is a novel antagonist of human constitutive androstane receptor. Mol Pharmacol. 74, 443–453

55. Seneviratne, M. S., Faccenda, D., De Biase, V., and Campanella, M. (2012) PK11195 inhibits mitophagy targeting the F1Fo-ATPsynthase in Bcl-2 knock-down cells. Current molecular medicine. 12, 476–482

56. Selvaraj, V., Morohaku, K., Koganti, P. P., Zhang, J., He, W., Quirk, S. M., and Stocco, D. M. (2020) Commentary: Amhr2-Cre-Mediated Global Tspo Knockout. Front. Endocrinol. 10.3389/fendo.2020.00472

57. Selvaraj, V., Tu, L. N., and Stocco, D. M. (2016) Crucial Role Reported for TSPO in Viability and Steroidogenesis is a Misconception. Commentary: Conditional Steroidogenic Cell-Targeted Deletion of TSPO Unveils a Crucial Role in Viability and Hormone-Dependent Steroid Formation. Front Endocrinol (Lausanne). 7, 91

58. Selvaraj, V., and Stocco, D. M. (2018) Letter to the Editor: Dubious Conclusions on TSPO Function. Endocrinology. 159, 2528–2529

59. Jamin, N., Neumann, J.-M., Ostuni, M. A., Vu, T. K. N., Yao, Z.-X., Murail, S., Robert, J.-C., Giatzakis, C., Papadopoulos, V., and Lacapère, J.-J. (2005) Characterization of the cholesterol recognition amino acid consensus sequence of the peripheral-type benzodiazepine receptor. Mol Endocrinol. 19, 588–594

60. Li, H., Yao, Z., Degenhardt, B., Teper, G., and Papadopoulos, V. (2001) Cholesterol binding at the cholesterol recognition/interaction amino acid consensus (CRAC) of the peripheral-type benzodiazepine receptor and inhibition of steroidogenesis by an HIV TAT-CRAC peptide. Proceedings of the National Academy of Sciences of the United States of America. 98, 1267–1272

61. Epand, R. M. (2006) Cholesterol and the interaction of proteins with membrane domains. Prog Lipid Res. 45, 279–294

62. Yang, G., Xu, H., Li, Z., and Li, F. (2014) Interactions of caveolin-1 scaffolding and intramembrane regions containing a CRAC motif with cholesterol in lipid bilayers. Biochimica et biophysica acta. 1838, 2588–2599

63. Thomas, J. L., Myers, R. P., and Strickler, R. C. (1989) Human placental 3 beta-hydroxy-5-ene-steroid dehydrogenase and steroid 5–4-ene-isomerase: purification from mitochondria and kinetic profiles, biophysical characterization of the purified mitochondrial and microsomal enzymes. J Steroid Biochem. 33, 209–217

64. Orme-Johnson, N. R., Light, D. R., White-Stevens, R. W., and Orme-Johnson, W. H. (1979) Steroid binding properties of beef adrenal cortical cytochrome P-450 which catalyzes the conversion of cholesterol into pregnenolone. Journal of Biological Chemistry. 254, 2103–2111

65. Ishikawa, M., Yoshitomi, T., Covey, D. F., Zorumski, C. F., and Izumi, Y. (2016) TSPO activation modulates the effects of high pressure in a rat ex vivo glaucoma model. Neuropharmacology. 111, 142–159

66. Koganti, P. P., Zhao, A. H., and Selvaraj, V. (2022) Exogenous cholesterol acquisition signaling in LH-responsive MA-10 Leydig cells and in adult mice. Journal of Endocrinology. 10.1530/JOE-22-0043

67. Staigmiller, R. B., Short, R. E., Bellows, R. A., and Carr, J. B. (1979) Effect of Nutrition on Response to Exogenous FSH in Beef Cattle. Journal of Animal Science. 48, 1182–1190

68. Turcu, A. F., Rege, J., Chomic, R., Liu, J., Nishimoto, H. K., Else, T., Moraitis, A. G., Palapattu, G. S., Rainey, W. E., and Auchus, R. J. (2015) Profiles of 21-Carbon Steroids in 21-hydroxylase Deficiency. J Clin Endocrinol Metab. 100, 2283–2290

69. Jaremko, L., Jaremko, M., Giller, K., Becker, S., and Zweckstetter, M. (2014) Structure of the mitochondrial translocator protein in complex with a diagnostic ligand. Science. 343, 1363–1366

70. Lomize, A. L., Todd, S. C., and Pogozheva, I. D. (2022) Spatial arrangement of proteins in planar and curved membranes by PPM 3.0. Protein Sci. 31, 209–220

